# Keap1 moderates the transcription of virus induced genes through G9a-GLP and NFкB p50 recruitment, and H3K9me2 deposition

**DOI:** 10.1101/2022.02.08.479619

**Authors:** Veronica Elizabeth Burns, Tom Klaus Kerppola

## Abstract

Cells must moderate transcription that is induced by virus infection to mitigate deleterious consequences of inflammation. We investigated the mechanisms whereby Keap1 moderates the transcription of genes that are induced by Sendai virus infection in mouse embryo fibroblasts (MEFs). Virus infection induced Keap1 to bind *Ifnb1, Tnf* and *Il6*, and reduced Keap1 binding at *Cdkn1a* and *Ccng1*. Keap1 was required for G9a and GLP to bind and to deposit H3K9me2 at these genes upon virus infection. Keap1 moderated the transcription of genes that were induced by virus infection in concert with G9a, GLP, and NFкB p50 recruitment. G9a-GLP lysine methyltransferase activity was required for Keap1 to moderate the transcription of virus induced genes. G9a-GLP inhibitors enhanced the transcription of virus induced genes, and they augmented Keap1 and NFкB p50 recruitment, in parallel with the inhibition of H3K9me2 deposition. The interdependent effects of Keap1 and G9a-GLP on transcription and on the recruitment of each other constitute a feedback circuit that moderates the transcription of virus induced genes. G9a-GLP inhibitors augmented Keap1 binding to different genes in virus infected and in uninfected MEFs, whereas they inhibited H3K9me2 deposition that was induced by virus infection selectively. G9a-GLP inhibitors stabilized Keap1 retention in permeabilized MEFs and augmented Keap1 binding to specific genes in parallel. Keap1 was required for NFкB p50 recruitment, and for the augmentation of NFкB binding by G9a-GLP inhibitors. Keap1 and the electrophile tBHQ attenuated virus induced gene transcription through independent mechanisms, and they regulated the recruitment of different NFкB subunits.

**Importance:** Excess and maladaptive immune responses to virus infections are a major contributing factor to the morbidity and mortality of COVID-19 and other diseases. Conversely, inadequate immune responses to vaccines and pathogens by individuals with suppressed immune function expose them to infections. Currently available drugs that enable therapeutic management of immune responses have low specificity and can blunt beneficial immune functions. The molecular mechanisms that moderate the transcription of genes that are induced by virus infection are incompletely understood. Characterization of the mechanisms whereby Keap1, G9a-GLP and NFкB p50 moderate virus induced gene transcription in mouse embryo fibroblasts represents the first step toward the identification of new targets for therapeutic agents that can modulate immune responsiveness.

## Introduction

Genes that are induced upon virus infection protect organisms from pathogenesis. Excess transcription of virus induced genes has deleterious effects. Elucidation of the factors and molecular mechanisms that moderate the transcription of virus induced genes is necessary to understand how homeostasis is maintained during viral infections.

*Keap1* (Kelch-like ECH-associated protein) deletion can cause spontaneous inflammation in several mouse organs as well as reduce experimentally induced inflammatory responses. Germline Keap1 deletion combined with conditional Nrf2 deletion in keratinocytes causes kidney inflammation that is associated with hydronephrosis and polyuria (1). Conditional Keap1 deletion in Tregs causes lung and liver inflammation (2). Conversely, conditional Keap1 deletions in Clara cells, in macrophages and granulocytes, in thymocytes, and in renal tubule cells reduce experimentally induced inflammatory responses (3-6). Since Keap1^fl^ alleles can be hypomorphic and their deletion is frequently inefficient, conditional Keap1 deletion may not produce loss of function phenotypes. The pro- and anti-inflammatory effects of *Keap1* deletions suggest that Keap1 influences immune functions through multiple mechanisms. To characterize the roles of Keap1 in immune regulation it is important to identify the cellular and molecular processes that mediate Keap1 effects on immune functions.

Keap1 depletion can enhance or reduce the induction of cytokine transcription in cultured cells. Keap1 depletion in RAW 264.7 and THP-1 macrophages/monocytes enhances *Il6* induction by LPS (7). Similarly, Keap1 depletion in human primary monocyte derived macrophages enhances inflammatory cytokine induction by *Mycobacterium avium* infection (8). These increases in cytokine induction correlate with increases in IKKα/β levels and phosphorylation (7, 8). In contrast, conditional *Keap1* deletion in mouse bone marrow derived macrophages reduces cytokine induction by LPS and IFNγ (9). This reduction of cytokine induction correlates with an increase in Nrf2. The contrasting effects of Keap1 depletion on cytokine induction in different macrophages could be due to indirect effects of Keap1 depletion, or they could be due to differences in the immune exposures of the macrophages prior to Keap1 depletion. To identify the direct and innate effects of Keap1 on cytokine transcription, it is important to determine how Keap1 regulates transcription in cells that have not been altered by prior immune exposures.

Fibroblasts respond rapidly to virus infection and serve as sentinel cells of the immune system (10). Lymphocytic choriomeningitis virus infection in mice induces immune response genes in the fibroblasts of many organs (11). Influenza virus infection in mice induces changes in lung fibroblast gene expression in regions of interstitial inflammation (12). Fibroblasts are a major constituent of the tumor microenvironment, which influences immune responses to tumor initiation, tumor growth, and immune checkpoint inhibitor therapeutics. Mouse embryo fibroblasts (MEFs) can be used to study mechanisms of gene regulation in naïve cells that have not been exposed to immune responses. Understanding of the mechanisms for regulation of the transcription of virus induced genes in MEFs provides a basis for the investigation of transcription regulation in other cells that control immune functions.

Sendai virus infection induces Keap1 to bind cytokine genes in MEFs (13). The levels of cytokine transcripts are higher in virus infected *Keap1-/-*MEFs than in MEFs with intact *Keap1*, suggesting that Keap1 moderates cytokine induction. To elucidate how Keap1 moderates the transcription of virus induced genes, it is necessary to determine the effects of Keap1 on transcription factor binding and on chromatin modifications, and their reciprocal effects on Keap1 binding to chromatin.

Keap1 binding to chromatin was discovered on *Drosophila* polytene chromosomes (14, 15). Keap1 binds to *Drosophila* genomic loci that encompass genes that are associated with innate immunity as well as other functions. It is important to compare the effects of virus infection on Keap1 binding to different classes of genes.

The G9a (encoded by *Ehmt2*) and GLP (encoded by *Ehmt1*) lysine methyltransferases catalyze histone H3 lysine 9 dimethylation (H3K9me2) (16). They have overlapping and interdependent activities (17). They are required for the differentiation and functions of both innate and adaptive immune cells (18-20). G9a and GLP are associated with the repression of cytokine transcription (21-23). It is important to determine the effects of virus infection on G9a and GLP binding to virus induced genes and the relationships between Keap1 and G9a-GLP recruitment.

Many pharmacological inhibitors of G9a and GLP have been developed (24-27). These compounds inhibit G9a and GLP lysine methyltransferase activities through different mechanisms and with different potencies. G9a and GLP can methylate substrates other than H3, and some functions of G9a and GLP do not require their lysine methyltransferase activities (18, 28). It is important to determine if H3K9me2 deposition or other G9a and GLP activities moderate regulatory protein binding to chromatin, or the transcription of virus induced genes.

NFкB p50 (encoded by *Nfkb1*) can both repress and activate cytokine transcription, whereas NFкB p65 (encoded by *RelA*) activates transcription (29-33). Conditional *Nfkb1* deletion in dendritic cells or in follicular B cells of bone marrow chimeric mice causes autoimmunity (34, 35). Heterozygous loss of function mutations in *NFKB1* are associated with both sporadic and familial common variable immunodeficiency (36, 37). Many of these patients present with autoinflammatory or autoimmune complications (38, 39). It is important to identify the differences in NFкB p50 and NFкB p65 regulation that can contribute to their distinct effects on immune functions, and to establish the effects of Keap1 on NFкB p50 and NFкB p65 binding to different genes in virus infected and in uninfected cells.

Keap1 binds electrophiles and represses electrophile response gene transcription through interaction with Nrf2. Some electrophiles reduce cytokine induction and autoinflammatory reactions (40). Dimethylfumarate is used to treat psoriasis and multiple sclerosis. The potential roles of Keap1 in electrophile effects on immunomodulatory gene transcription need to be evaluated.

We tested the hypothesis that Sendai virus infection induces Keap1 to bind specific genes, and that Keap1 moderates the transcription of those genes. We examined the relationships between Keap1 and G9a-GLP recruitment and H3K9me2 deposition at virus induced genes, and the interdependence of their effects on virus induced transcription. We investigated the roles of lysine methyltransferase activity in G9a-GLP effects on transcription and on recruitment of other proteins at virus induced genes by using several structurally dissimilar G9a-GLP inhibitors. We compared the effects of Keap1 on NFкB p50 and NFкB p65 binding to different genes and to proximal and distal elements. We also compared the effects of Keap1 and of electrophiles on transcription and on NFкB subunit recruitment at virus induced genes. We found that Keap1, G9a, GLP and NFкB p50 regulated recruitment of each other, and moderated transcription at virus induced genes through H3K9me2 deposition.

## Results

We compared the effects of Keap1 on the transcription of genes that were induced by Sendai virus infection and of uninduced genes in MEFs. We measured the transcript levels in MEFs with intact *Keap1* and in MEFs with *Keap1-/-* deletions (Fig. 1A). To avoid the indirect effects of *Keap1-/-* deletions that result from constitutive Nrf2 activation, we focused on the effects of *Keap1-/-* deletions in MEFs that also carried *Nrf2-/-* deletions (*Keap1-/-* MEFs). We measured transcript levels at different times after virus infection to characterize the effects of Keap1 on the primary transcriptional response.

**Figure 1.**
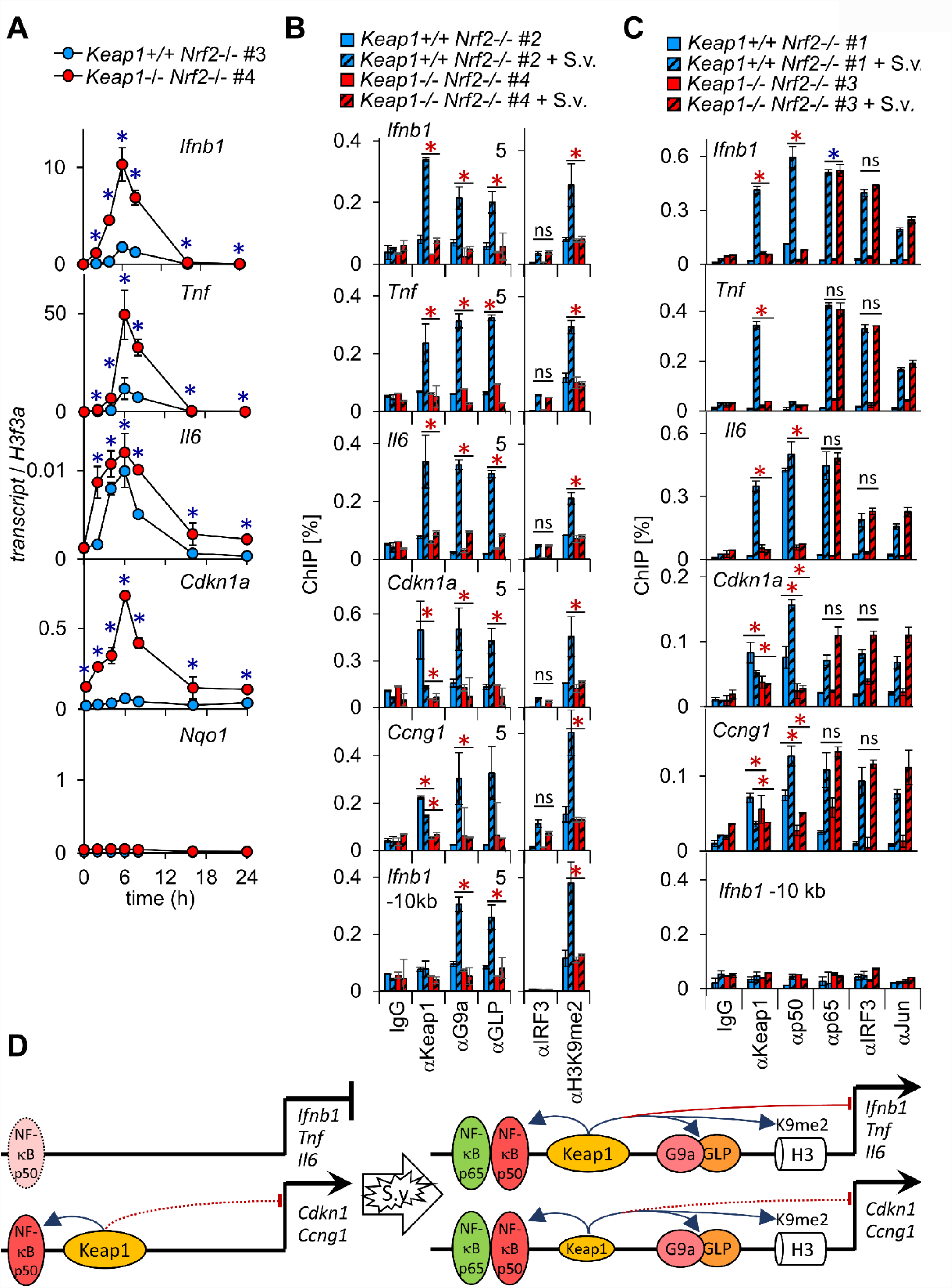
Virus infection induces higher cytokine transcript levels in *Keap1-/-* MEFs than in MEFs with intact *Keap1*, and has opposite effects on Keap1 binding at cytokine and at cell cycle genes; Keap1 is required for G9a, GLP and NFкB p50 recruitment, and for H3K9me2 deposition. (A) Virus infection induces higher levels of cytokine transcription in *Keap1-/-* MEFs than in MEFs with intact *Keap1*. MEFs with intact *Keap1* (blue) and MEFs with *Keap1-/-* deletions (red), each with *Nrf2-/-* deletions, were infected with Sendai virus. The levels of the transcripts indicated in the graphs were measured by RT-qPCR at the times after virus infection indicated on the bottom graph. The line graphs show the results (mean±2*SD) of a representative experiment in which *Keap1+/+ Nrf2-/-* #3 and *Keap1-/- Nrf2-/-* #4 MEFs were compared. The # after each genotype reflects a MEF population that was isolated from an independent embryo. The reproducibility of Keap1 effects on transcript levels were evaluated in 3-7 independent sets of MEFs by performing two-factor ANOVA analyses (* p<0.0005). (B) Virus infection has opposite effects on Keap1 binding to cytokine versus cell cycle genes, and Keap1 is required for G9a and GLP to bind and to deposit H3K9me2 upon virus infection. MEFs with intact Keap1 (blue bars) and MEFs with *Keap1-/-* deletions (red bars), each with *Nrf2-/-* deletions, were infected with mock (solid bars) or Sendai virus (S.v., striped bars). The levels of Keap1, G9a, GLP and IRF3 binding, and of H3K9me2 were measured 6 hours after infection at the genes indicated in the graphs using the antibodies indicated. The bar graphs show the results (mean±2*SD) of a representative experiment in which *Keap1+/+ Nrf2-/-* #2 and *Keap1-/- Nrf2-/-* #4 MEFs were compared. The reproducibility of Keap1 effects on G9a, GLP and IRF3 binding, and on H3K9me2 were evaluated in 2-5 independent sets of MEFs by two-factor ANOVA analyses (* p<0.001, blue – increase, red – decrease). (C) Keap1 is required for NFкB p50 to bind both virus induced and uninduced genes. MEFs with intact Keap1 (blue bars) and MEFs with *Keap1-/-* deletions (red bars), each with *Nrf2-/-* deletions, were infected with mock (solid bars) or Sendai virus (S.v. striped bars). Chromatin was isolated. The levels of NFкB p50, NFкB p65, IRF3 and cJun binding were measured 6 hours after infection at the genes indicated in the graphs using the antibodies indicated at the bottom. The bar graphs show the results (mean±2*SD) of a representative experiment in which *Keap1+/+ Nrf2-/-* #1 and *Keap1-/- Nrf2-/-* #3 MEFs were compared. The reproducibility of Keap1 effects on NFкB p50 and NFкB p65 binding were evaluated in 4-6 independent sets of MEFs by two-factor ANOVA analyses (* p<0.001, blue – increase, red – decrease). (D) The diagrams compare effects of virus infection on Keap1 binding, and Keap1 effects on G9a, GLP and NFкB p50 binding, on H3K9me2 deposition, and on transcription at cytokine *versus* cell cycle associated genes. The blue arrows indicate effects that are required for binding or for deposition. The red arcs and bars indicate effects that moderate transcription. Dotted lines and ovals indicate differences between different genes and MEFs.

Virus infection induced *Ifnb1, Tnf* and *Il6* transcription more rapidly and to higher levels in *Keap1-/-* MEFs than in MEFs with intact *Keap1* (Fig. 1A). The higher transcript levels were observed beginning at 2 h after virus infection. The peak *Ifnb1, Tnf* and *Il6* transcript levels were 3-to 8-fold higher on average in 7 different sets of *Keap1-/-* MEFs relative to MEFs with intact *Keap1*. Keap1 reduced the levels of virus induced transcripts at the earliest times after infection that were tested by mechanisms that were independent of Nrf2, consistent with direct moderation of the transcription of virus induced genes by Keap1.

We examined the effects of Keap1 on cell cycle associated gene transcription because Keap1 interacts with DNA replication factors, and because the stability of Keap1 retention in the nucleus varies at different stages of the cell cycle (41). The basal levels of *Cdkn1a* and *Ccng1* transcripts were higher in *Keap1-/-* MEFs than in MEFs with intact *Keap1* (Fig. 1A). Virus infection induced *Cdkn1a* transcription in two of the three independent *Keap1-/-* MEFs that were tested. In contrast, virus infection did not induce *Cdkn1a* transcription in MEFs with intact *Keap1*. Latent virus induced transcription of *Cdkn1a* was suppressed in MEFs with intact *Keap1*.

There was no difference in the accumulation of Sendai virus M gene transcripts between *Keap1-/-* MEFs and MEFs with intact *Keap1* at early times after virus infection (Fig. 2A). This indicates that Keap1 did not affect the efficiencies of virus infection or replication. The small effects of virus infection on *Ccng1, Gapdh* and *Nqo1* transcription did not differ between *Keap1-/-* MEFs and MEFs with intact *Keap1* (Fig. 1A, 2A, 3B). Keap1 reduced the levels of virus induced transcripts selectively.

**Figure 2.**
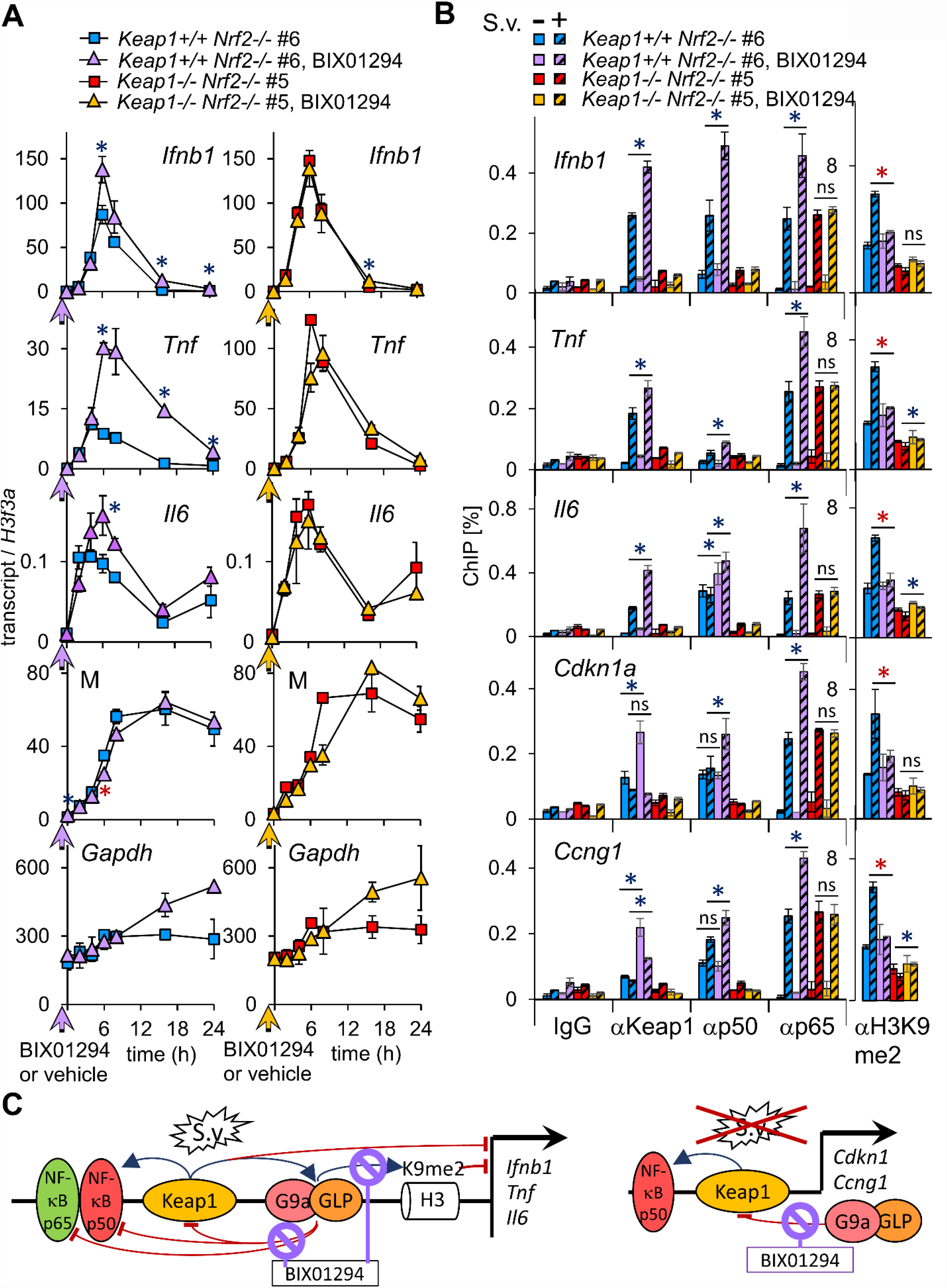
Inhibition of G9a-GLP enhances the transcription of virus induced genes in MEFs with intact Keap1, and augments Keap1 binding to different genes in uninfected and in virus infected MEFs. (A) BIX01294 enhances the transcription of virus induced genes in MEFs with intact *Keap1* selectively. MEFs with intact Keap1 (left graphs) and MEFs with *Keap1-/-* deletions (right graphs), each with *Nrf2-/-* deletions, were cultured with 20 µM BIX01294 or vehicle starting an hour before virus infection (arrowhead on time axis). The levels of the transcripts indicated in the graphs were measured at the times after virus infection indicated at the bottom. The line graphs show the results of a representative experiment in which *Keap1+/+ Nrf2-/-* #6 and *Keap1-/- Nrf2-/-* #5 MEFs were compared. The reproducibility of BIX01294 effects on transcription were evaluated in 2-4 independent sets of MEFs by two-factor ANOVA analyses (* p<0.001, blue – increase, red – decrease). (B) BIX01294 augments Keap1 binding to different genes in virus infected and in uninfected MEFs. MEFs with intact *Keap1* and MEFs with *Keap1-/-* deletions, each with *Nrf2-/-* deletions, were cultured with 20 µM BIX01294 or vehicle for an hour before mock (solid bars) or virus (striped bars) infection. The levels of Keap1, NFкB p50 and NFкB p65 binding, and of H3K9me2 were measured 6 hours after infection at the genes indicated in the graphs using the antibodies indicated. The bar graphs show the results of a representative experiment in which *Keap1+/+ Nrf2-/-* #6 and *Keap1-/- Nrf2-/-* #5 MEFs were compared. The reproducibility of BIX01294 effects on Keap1, NFкB p50 and NFкB p65 binding, and on H3K9me2 were evaluated in 2-5 independent sets of MEFs by two-factor ANOVA analyses (* p<0.005, blue – increase, red – decrease). (C) The diagrams compare BIX01294 effects on Keap1 and NFкB p50 binding, on H3K9me2 deposition, and on transcription at cytokine genes in virus infected MEFs and at cell cycle associated genes in uninfected MEFs. The blue arrows indicate effects that are required for binding or for deposition. The red arcs and bars indicate effects that inhibit or moderate binding, deposition, or transcription.

We compared the effects of Keap1 on virus induced gene (*Ifnb1, Tnf* and *Il6*) transcription and on electrophile response gene (*Nqo1*) transcription in MEFs with intact *Nrf2* and in MEFs with *Nrf2-/-* deletions. The *Nqo1* transcript level was 50-fold higher in *Keap1-/-* MEFs with intact *Nrf2* than in MEFs with intact *Keap1* and *Nrf2*. There was little difference in *Nqo1* transcript levels between *Keap1-/- Nrf2-/-* MEFs and *Nrf2-/-* MEFs (Fig. 1). In contrast, virus infection induced higher *Ifnb1, Tnf* and *Il6* transcript levels in *Keap1-/- Nrf2-/-* MEFs than in *Nrf2-/-* MEFs (Fig. 1). The smaller effects of *Keap1-/-* deletions on *Ifnb1, Tnf* and *Il6* transcription in MEFs with intact Nrf2 than in MEFs that also contained *Nrf2-/-* deletions could be attributable to the effects of *Keap1-/-* deletions on the growth rate and NFкB activation in MEFs with intact *Nrf2*, but not in MEFs with *Nrf2-/-* deletions (13). Keap1 moderated the transcription of virus induced genes by mechanisms that did not require Nrf2, and that were distinct from Keap1 regulation of electrophile response gene transcription.

### Virus infection induces Keap1 to bind cytokine genes and reduces Keap1 binding at cell cycle genes

We compared the effects of virus infection on Keap1 binding to the promoter regions of cytokine and cell cycle associated genes. Virus infection induced Keap1 to bind *Ifnb1, Tnf* and *Il6* in the 7 independent MEFs with intact *Keap1* that were tested, and not in *Keap1-/-* MEFs (Fig. 1B, 1C). The αKeap1 ChIP signals at these genes were 11- to 14-fold higher, on average, in virus infected MEFs than in uninfected MEFs. We determined that the ChIP signals reflected Keap1 binding to virus induced genes by using antibodies that recognized different epitopes in Keap1 and by employing several independent criteria to establish the validity of the ChIP signals. Since Keap1 bound to these genes upon virus infection, and since the levels of these transcripts were higher in *Keap1-/-* MEFs, we inferred that Keap1 moderated their transcription by binding to the genes upon virus infection.

Keap1 bound to *Cdkn1a* and *Ccng1* in uninfected MEFs. Virus infection reduced Keap1 binding to these genes in the 4 independent MEFs that were tested (Fig. 1B, 1C). The αKeap1 ChIP signals at *Cdkn1a* and *Ccng1* were 21% and 23% lower, on average, in virus infected MEFs than in uninfected MEFs. The differences in the effects of virus infection on Keap1 binding to *Ifnb1, Tnf* and *Il6* versus *Cdkn1a* and *Ccng1* suggest that virus infection regulated Keap1 binding to different genes by selective mechanisms.

### Keap1 is necessary but not sufficient for G9a and GLP to bind and to deposit H3K9me

G9a and GLP bind to cytokine genes in virus infected MEFs with intact Keap1, but not in *Keap1-/-* MEFs (13). We compared G9a and GLP binding at different classes of genes in virus infected and in uninfected MEFs with intact Keap1 and with *Keap1-/-* deletions to investigate the relationships among Keap1, G9a and GLP binding. Virus infection induced G9a and GLP to bind *Ifnb1, Tnf* and *Il6* in the 3 independent MEFs with intact *Keap1* that were tested (Fig. 1B). Virus infection also induced G9a and GLP to bind *Cdkn1a* and *Ccng1*, and 10 kb upstream of *Ifnb1* in MEFs with intact *Keap1*. Virus infection did not induce G9a or GLP to bind the genes that were examined in *Keap1-/-* MEFs. The αG9a and αGLP ChIP signals at these genes were 3- to 7-fold higher on average in virus infected MEFs with intact *Keap1* than in *Keap1-/-* MEFs. Since Keap1 was required for both G9a and GLP binding, and since both G9a and GLP have lysine methyltransferase activity and can form heterodimers, we refer to these complexes collectively as G9a-GLP. Keap1 was required for G9a-GLP recruitment both to virus induced genes and to genes that are not induced by virus infection in virus infected MEFs, but Keap1 binding to chromatin was not sufficient for G9a-GLP recruitment in uninfected MEFs.

Since G9a-GLP catalyze histone H3K9 dimethylation, we examined the effects of virus infection on H3K9me2 at different genes in MEFs with intact Keap1 and in MEFs with *Keap1-/-* deletions. Virus infection induced H3K9me2 at the genes that were examined in the 6 independent MEFs with intact Keap1 that were tested (Fig. 1B). By contrast, virus infection did not increase H3K9me2 at these genes in *Keap1-/-* MEFs. It is likely that the lack of virus induced H3K9me2 deposition in *Keap1-/-* MEFs was due to the lack of virus induced G9a or GLP binding. Significantly, the basal H3K9me2 levels were the same in *Keap1-/-* MEFs and in MEFs with intact *Keap1*, indicating that Keap1 affected virus induced H3K9me2 deposition selectively. Virus infection reduced H3K27me3 at these genes, indicating that the increase in the H3K9me2 signal upon virus infection was not due to an increase in chromatin accessibility (Fig. 3A). Additionally, the reduction in H3K27me3 was equivalent in *Keap1-/-* MEFs and in MEFs with intact *Keap1*, indicating that Keap1 affected H3K9me2 deposition selectively. Keap1 was required for G9a-GLP to bind and to deposit H3K9me2, and for the moderation of virus induced gene transcription.

**Figure 3.**
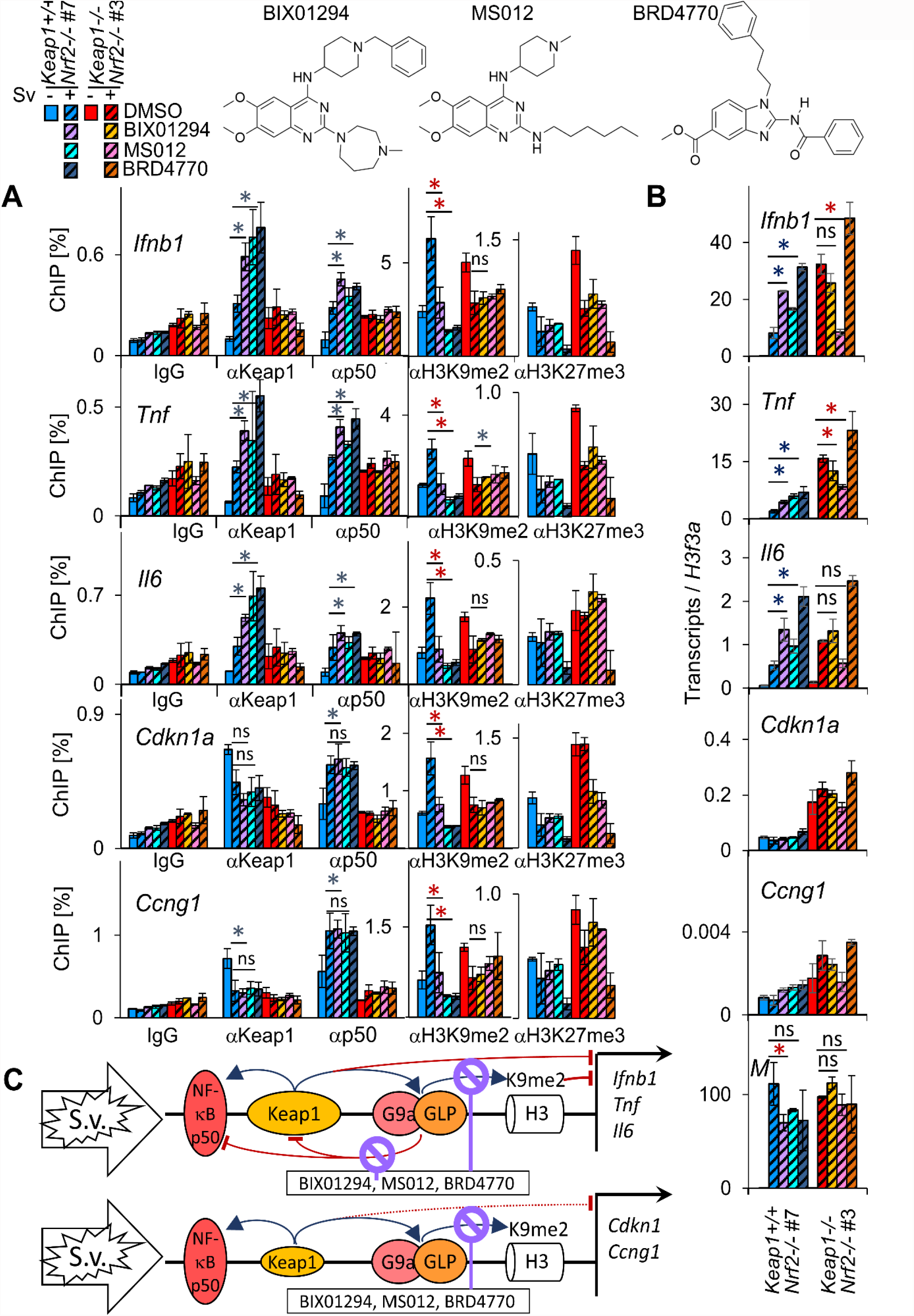
Structurally dissimilar G9a-GLP inhibitors augment Keap1 and NFкB p50 binding to virus induced genes, and enhance the transcription of virus induced genes in MEFs with intact *Keap1*. (A) Different G9a-GLP inhibitors have parallel effects on Keap1 and NFкB p50 binding to virus induced genes in MEFs with intact *Keap1*. MEFs with intact Keap1 (cool colors) and MEFs with *Keap1-/-* deletions (warm colors), each with *Nrf2-/-* deletions, were cultured with 20 µM BIX01294 starting an hour before infection, or with 1 µM MS012, 20 µM BRD4770 or vehicle starting 48 hours before infection. The levels of Keap1 and NFкB p50 binding, and H3K9me2 and H3K27me3 were measured 6 hours after mock (solid bars) or virus (striped bars) infection at the genes that are indicated in the graphs using the antibodies indicated at the bottom. The bar graphs show the results of a representative experiment in which *Keap1+/+ Nrf2-/-* #7 and *Keap1-/- Nrf2-/-* #3 MEFs were compared. The MEFs that were analyzed in panels A and B were cultured in parallel, and the legend at the top of the figure applies to both panels. The reproducibility of BIX01294 and MS012 effects on Keap1 and NFкB p50 binding, and on H3K9me2 were evaluated in 2-4 independent sets of MEFs by two-factor ANOVA analyses (* p<0.005, blue – increase, red – decrease). (B) Several G9a-GLP inhibitors have distinct effects on virus induced gene transcription in MEFs with intact Keap1 *versus* MEFs with *Keap1-/-* deletions. MEFs with intact Keap1 (cool colors) and MEFs with *Keap1-/-* deletions (warm colors), each with *Nrf2-/-* deletions, were cultured with 20 µM BIX01294 starting an hour before infection, or with 1 µM MS012, 20 µM BRD4770 or vehicle starting 48 hours before infection. The levels of the transcripts indicated in the graphs were measured 6 hours after mock (solid bars) or virus (striped bars) infection. The graphs show the results of a representative experiment in which *Keap1+/+ Nrf2-/-* #7 and *Keap1-/- Nrf2-/-* #3 MEFs were compared. The reproducibility of BIX01294 and MS012 effects on transcription were evaluated in 2-4 independent sets of MEFs by two-factor ANOVA analyses (* p<0.001, blue – increase, red – decrease). (C) The diagrams compare effects of G9a-GLP inhibitors on Keap1 and NFкB p50 binding, on H3K9me2 deposition, and on transcription at cytokine *versus* cell cycle associated genes in virus infected MEFs. The blue arrows indicate effects that are required for binding or for deposition. The red arcs and bars indicate effects that inhibit or moderate binding, deposition, or transcription.

### Keap1 as well as virus infection have distinct effects on NFкB p50 versus NFкB p65 binding

NFкB p50 binds to cytokine genes in virus infected MEFs with intact Keap1, but not in *Keap1-/-* MEFs (13). We compared the effects of Keap1 on the binding of different NFкB subunits and of other transcription factors to different genes in virus infected and in uninfected MEFs. Virus infection induced NFкB p50 to bind *Ifnb1* in the 6 independent MEFs with intact *Keap1* that were tested (Fig. 1C). Virus infection did not induce NFкB p50 to bind *Ifnb1* in 5 independent *Keap1-/-* MEFs. Virus infection induced different levels of NFкB p50 binding to *Tnf* in different experiments for reasons that are not known (Fig. 1C, 2B, 3A, 6B). Nevertheless, virus infection induced NFкB p50 to bind *Tnf* in some experiments with each of the 6 independent MEFs with intact *Keap1* that were tested. Virus infection did not induce NFкB p50 to bind *Tnf* in any experiments with the 5 independent *Keap1-/-* MEFs. NFкB p50 bound to *Il6* in 2 of the 6 uninfected MEFs with intact *Keap1*, and in all 6 MEFs with intact *Keap1* upon virus infection (Fig. 1C, 2B, 3A). NFкB p50 did not bind to *Il6* in the 5 independent *Keap1-/-* MEFs in the absence or in the presence of virus. Consequently, Keap1 was required for NFкB p50 to bind *Ifnb1, Tnf* and *Il6* upon virus infection.

NFкB p50 bound to *Cdkn1a* and *Ccng1* in the absence of virus infection in the 5 independent MEFs with intact *Keap1* that were tested (Fig. 1B, 2B.3A). Virus infection augmented NFкB p50 binding to *Cdkn1a* and *Ccng1* in MEFs with intact *Keap1*. NFкB p50 did not bind to *Cdkn1a* or *Ccng1* in *Keap1-/-* MEFs in the absence or in the presence of virus infection. *Keap1* was required for NFкB p50 to bind *Cdkn1a* and *Ccng1* in uninfected and in virus infected MEFs.

NFкB p65 bound to *Cdkn1a* and *Ccng1* only in virus infected MEFs, whereas NFкB p50 bound to these genes also in uninfected MEFs. Virus infection induced equal or higher levels of NFкB p65 binding in *Keap1-/-* MEFs than in MEFs with intact Keap1 at all genes that were examined, whereas virus infection induced little or no NFкB p50 binding in *Keap1-/-* MEFs at any of the genes. Thus, both virus infection and Keap1 had distinct effects on NFкB p50 *versus* NFкB p65 binding at the genes that were tested.

Virus infection induced equivalent levels of IRF3 as well as cJun binding in *Keap1-/-* MEFs and in MEFs with intact Keap1 at the genes that were tested (Fig. 1B, 1C). Thus, the requirement for Keap1 was unique to G9a-GLP and NFкB p50 recruitment among the chromatin binding proteins that were tested.

### G9a-GLP inhibition enhances virus induced gene transcription in MEFs with intact *Keap1*

Inhibitors of G9a and GLP lysine methyltransferase activities enhance cytokine transcription and augment Keap1 binding at cytokine genes (13, 23). We measured the effects of G9a-GLP inhibitors on virus induced gene transcription in MEFs with intact Keap1 and MEFs with *Keap1-/-* deletions to investigate the relationships between Keap1 and G9a-GLP in the moderation of virus induced gene transcription. The G9a-GLP inhibitor BIX01294 (42) was added to the MEFs one hour before virus infection to focus on the direct effects of G9a-GLP inhibition. BIX01294 enhanced *Ifnb1, Tnf*, and *Il6* transcription in the 4 independent MEFs with intact *Keap1* that were tested (Fig. 2A, left column). The peak *Ifnb1, Tnf* and *Il6* transcript levels were 2.3- to 2.4-fold higher on average in the MEFs with intact *Keap1* that were cultured with BIX01294 than in the same MEFs that were cultured with vehicle. In contrast, BIX01294 did not increase the peak *Ifnb1, Tnf*, or *Il6* transcript levels in *Keap1-/-* MEFs (Fig. 2A, right column). BIX01294 and *Keap1-/-* deletions separately had quantitatively equivalent effects on *Ifnb1* as well as *Il6* transcript levels. Since Keap1 moderated the transcription of virus induced genes, and since BIX01294 enhanced the transcription of virus induced genes selectively in MEFs with intact *Keap1*, we inferred that BIX01294 counteracted the ability of Keap1 to moderate the transcription of virus induced genes.

BIX01294 did not increase viral M gene transcript accumulation in MEFs with intact Keap1 or in *Keap1-/-* MEFs (Fig. 2A). BIX01294 had little or no effect on the levels of *Gapdh, Cdkn1a* or *Ccng1* transcripts within 6 h after virus infection (Fig. 2A, 3B). BIX01294 enhanced virus induced gene transcription rapidly and selectively in MEFs with intact *Keap1*.

### G9a-GLP inhibition augments Keap1 binding to different genes in uninfected and in virus infected MEFs

We compared the effects of G9a-GLP lysine methyltransferase inhibitors on Keap1 binding to virus induced and to uninduced genes in virus infected and in uninfected MEFs. BIX01294 augmented Keap1 binding to *Ifnb1, Tnf* and *Il6* upon virus infection in the 5 independent MEFs with intact Keap1 that were tested (Fig. 2B, 3B). The αKeap1 ChIP signals at these genes were 1.9- to 2.1-fold higher, on average, in virus infected MEFs that were cultured with BIX01294 than in the same MEFs that were cultured with vehicle. BIX01294 augmented the low αKeap1 ChIP signals at these genes in uninfected MEFs, but these signals remained near the IgG background. The augmentation of Keap1 binding by BIX01294 was detected by antibodies that were raised against different regions of Keap1.

BIX01294 augmented Keap1 binding to *Cdkn1a* and *Ccng1* in uninfected MEFs. The αKeap1 ChIP signals at *Cdkn1a* and *Ccng1* were 1.9 and 2.8 higher, on average, in uninfected MEFs that were cultured with BIX01294, than in the same MEFs that were cultured with vehicle (Fig. 2B).

Whereas virus infection and BIX01294 cooperated to augment Keap1 binding to *Ifnb1, Tnf* and *Il6*, virus infection counteracted the effects of BIX01294 on Keap1 binding to *Cdkn1a* and *Ccng1*. The effects of BIX01294 on Keap1 binding to *Cdkn1a* and *Ccng1* in different virus infected MEFs depended on the efficiency of viral inhibition of Keap1 binding at each gene (Fig. 2B, 3A). Consequently, BIX01294 augmented Keap1 binding to *Ifnb1, Tnf* and *Il6* mainly in virus infected MEFs, whereas BIX01294 augmented Keap1 binding to *Cdkn1a* and *Ccng1* more efficiently in uninfected MEFs.

### G9a-GLP inhibition augments virus induced NFкB binding in MEFs with intact *Keap1*

We compared BIX01294 effects on NFкB p50 and NFкB p65 binding to different genes in MEFs with intact *Keap1* and in *Keap1-/-* MEFs. BIX01294 augmented viral induction of NFкB p50 binding to *Ifnb1, Tnf* and *Il6* in the 5 independent MEFs with intact *Keap1* that were tested (Fig. 2B, 3A). The αp50 ChIP signals were 1.6- to 1.9-fold higher, on average, in virus infected MEFs with intact *Keap1* that were cultured with BIX01294 than they were in the same MEFs that were cultured with vehicle. The effects of BIX01294 on NFкB p50 binding to *Cdkn1a* and *Ccng1* correlated with BIX01294 effects on Keap1 binding in the same MEFs (Fig. 2B, 3A). BIX01294 did not affect the low αp50 ChIP signals at the genes examined in *Keap1-/-* MEFs. Since BIX01294 had parallel effects on NFкB p50 and Keap1 binding to the genes that were examined, and since Keap1 was essential for NFкB p50 binding, it is likely that the effects of BIX01294 on NFкB p50 binding were mediated by BIX01294 effects on Keap1 binding to these genes.

BIX01294 augmented NFкB p65 binding to the genes that were examined in virus infected MEFs with intact *Keap1* (Fig. 2B). In contrast, BIX01294 did not augment viral induction of NFкB p65 binding to these genes in *Keap1-/-* MEFs. Thus, Keap1 was specifically required for BIX01294 to augment NFкB p65 binding to these genes, whereas Keap1 was not required for viral induction of NFкB p65 binding. The requirement for Keap1 in the augmentation of NFкB p65 binding by BIX01294 suggests that G9a-GLP moderates NFкB p65 binding only when G9a-GLP binding to the same genes is facilitated by Keap1.

BIX01294 inhibited virus induced H3K9me2 deposition in MEFs with intact *Keap1* (Fig. 2B, 3A). In contrast, BIX01294 had no effect on basal H3K9me2 in uninfected MEFs or in *Keap1-/-* MEFs. BIX01294 did not affect the αH3 or αH3K27me3 ChIP signals in any of the MEFs (Fig. 3A). The effects of BIX01294 on Keap1 and NFкB p50 binding and on transcription correlated with BIX01294 effects on H3K9me2 deposition at virus induced genes (*Ifnb1, Tnf* and *Il6*), but not at uninduced genes (*Cdkn1a* and *Ccng1*).

### Structurally dissimilar G9a-GLP inhibitors augment Keap1 and NFкB p50 binding to virus induced genes

We compared the effects of five G9a-GLP inhibitors on Keap1 and on NFкB p50 binding to virus induced genes. MS012 interacts with the peptide binding sites of G9a and GLP (27). MS012 augmented Keap1 and NFкB p50 binding to *Ifnb1, Tnf* and *Il6* in the three independent MEFs with intact *Keap1* that were tested (Fig. 3A). The αKeap1 ChIP signals at these genes were 2.3- to 2.7-fold higher, and the αp50 signals were 1.9- to 2.0-fold higher, in virus infected MEFs that were cultured with MS012 than in the same MEFs that were cultured with vehicle. MS012 had parallel effects on Keap1 and on NFкB p50 binding to different genes in virus infected MEFs (Fig. 3A). MS012 did not affect the low αp50 ChIP signals in *Keap1-/-* MEFs. MS012 inhibited virus induced H3K9me2 deposition in MEFs with intact *Keap1* (Fig. 3A). MS012 augmented Keap1 and NFкB p50 binding and inhibited H3K9me2 deposition at a lower concentration than BIX01294. The parallel effects of BIX01294 and MS012 both on Keap1 and NFкB p50 binding, and on H3K9me2 deposition indicate that these compounds augmented Keap1 and NFкB p50 binding by inhibiting G9a-GLP lysine methyltransferase activities at virus induced genes.

BRD4770 inhibits lysine methyltransferases by competition with S-adenosyl methionine and its overall structure differs from those of the other G9a-GLP inhibitors that were tested (26). BRD4770 augmented Keap1 and NFкB p50 binding and inhibited H3K9me2 deposition at virus induced genes in MEFs with intact *Keap1* (Fig. 3A). BRD4770 reduced H3K27me3 at these genes both in MEFs with intact Keap1 and in *Keap1-/-* MEFs, suggesting that it inhibited lysine methyltransferases other that G9a-GLP independently of Keap1. UNC0638 and UNC0642 augmented Keap1 and NFкB p50 binding and inhibited H3K9me2 deposition at virus induced genes with different efficiencies in different experiments. These differences correlated with differences in their inhibition of H3K9me2 deposition in different MEFs (Fig. 4). Other compounds, including dexamethasone and *tert*-butylhydroquinone (tBHQ) did not augment Keap1 or NFкB p50 binding to virus induced genes, and had no effect on H3K9me2 deposition (Fig. 6B). Thus, structurally dissimilar G9a-GLP lysine methyltransferase inhibitors augmented Keap1 and NFкB p50 binding, and inhibited H3K9me2 deposition at virus induced genes.

**Figure 4.**
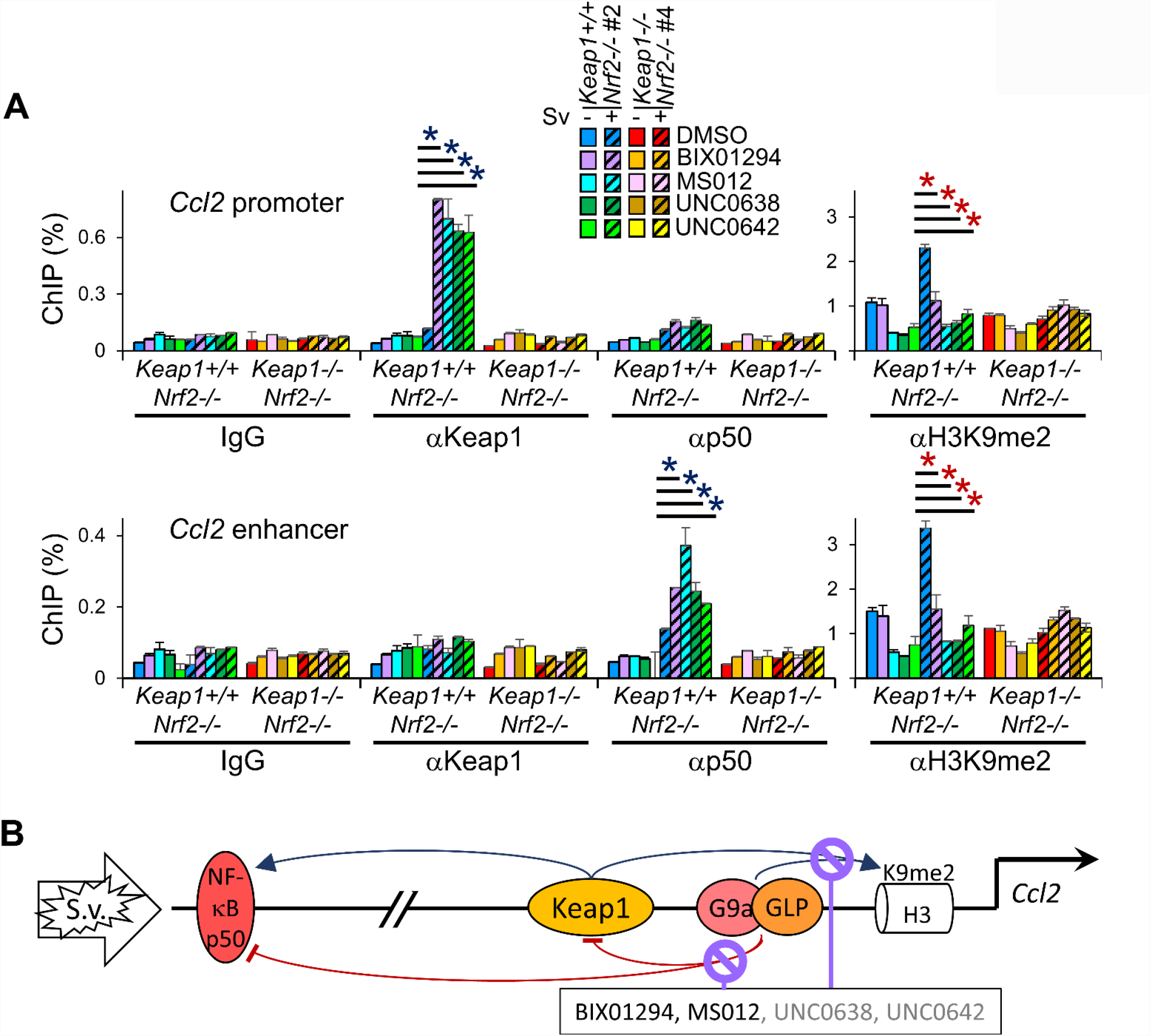
G9a-GLP inhibitors augment Keap1 binding to the *Ccl2* promoter and NFкB p50 binding to the *Ccl2* enhancer in virus infected MEFs. (A) Keap1 binds to the *Ccl2* promoter and is required for NFкB p50 to bind the *Ccl2* enhancer and for H3K9me2 deposition. MEFs with intact Keap1 (cool colors) and MEFs with *Keap1-/-* deletions (warm colors), each with *Nrf2-/-* deletions, were cultured for one hour with 20 µM BIX01294, or for 48 hours with 1 µM MS012, 10 µM UNC0638, 10 µM UNC0642, or vehicle before infection. The levels of Keap1 and NFкB p50 binding, and of H3K9me2 were measured 6 h after mock (solid bars) or virus (striped bars) infection at the *Ccl2* promoter (upper graph) and at the *Ccl2* enhancer (lower graph) using the antibodies indicated. The graphs show the results of a representative experiment in which *Keap1+/+ Nrf2-/-* #2 and *Keap1-/- Nrf2-/-* #4 MEFs were compared. The reproducibility of G9a-GLP inhibitor effects on Keap1 and NFкB p50 binding, and on H3K9me2 were evaluated in 2 experiments by two-factor ANOVA analyses (* p<0.005, blue – increase, red – decrease). (B) The diagram depicts effects of virus infection and G9a-GLP inhibitors on Keap1 and NFкB p50 binding and on H3K9me2 deposition at the *Ccl2* enhancer and promoter in virus infected MEFs. The blue arrows indicate effects that are required for binding or for deposition. The red arcs and bars indicate effects that inhibit or moderate binding or deposition. Compounds that had different effects in different experiments are indicated in grey type.

### Compounds that inhibit G9a-GLP through different mechanisms enhance virus induced gene transcription in MEFs with intact *Keap1*

We compared the effects of structurally dissimilar G9a-GLP inhibitors on virus induced gene transcription. BIX01294, MS012 and BRD4770 enhanced *Ifnb1, Tnf* and *Il6* transcription in virus infected MEFs with intact *Keap1* (Fig. 3B). MS012 enhanced these transcript levels 2.1- to 2.3-fold on average in the two independent MEFs with intact *Keap1* that were tested. BRD4770 enhanced transcription in *Keap1-/-* MEFs, whereas UNC0638 and UNC0642 inhibited transcription in *Keap1-/-* MEFs (Fig. 3B). These effects could be due to the inhibition of other lysine methyltransferases, as suggested by the reduction of H3K27me3 by BRD4770 (Fig. 3A). BIX01294, MS012 and BRD4770 had little effect on *Cdkn1a* or *Ccng1* transcription and did not increase viral M gene transcript accumulation. The preferential enhancement of virus induced gene transcription by BIX01294, MS012 and BRD4770 in MEFs with intact *Keap1* suggests that G9a-GLP inhibition counteracted the ability of Keap1 to moderate the transcription virus induced genes.

### Keap1 is required for NFкB p50 to bind the Ccl2 enhancer

We investigated if Keap1 was required for NFкB p50 to bind the *Ccl2* enhancer, which is located at a distance from the *Ccl2* promoter. Virus infection induced Keap1 to bind the *Ccl2* promoter and NFкB p50 to bind the *Ccl2* enhancer in MEFs with intact *Keap1* (Fig. 4). A low level of NFкB p50 bound to the *Ccl2* promoter. Virus infection did not induce NFкB p50 to bind the *Ccl2* enhancer or promoter in *Keap1-/-* MEFs. Keap1 was required for NFкB p50 to bind the *Ccl2* enhancer, though Keap1 binding was detected mainly at the *Ccl2* promoter.

We examined the effects of G9a-GLP inhibitors on Keap1 and NFкB p50 binding to the *Ccl2* promoter and enhancer. The G9a-GLP inhibitors augmented Keap1 binding to the *Ccl2* promoter and NFкB p50 binding to the *Ccl2* enhancer upon virus infection (Fig. 4). The G9a-GLP inhibitors augmented NFкB p50 binding to the *Ccl2* enhancer in virus infected MEFs with intact *Keap1*, and not in *Keap1-/-* MEFs. The augmentation of Keap1 and NFкB p50 binding to the *Ccl2* promoter and enhancer by different G9a-GLP inhibitors correlated with the inhibition of virus induced H3K9me2 deposition by these compounds (Fig. 4).

### G9a-GLP inhibitors strengthen Keap1 retention in permeabilized MEFs

To determine if G9a-GLP inhibitors affected the stability of Keap1 interactions in cells, we compared Keap1 retention in MEFs that were cultured with BIX01294 or with vehicle, followed by incubation with detergents. Stronger Keap1 binding to chromatin is predicted to increase Keap1 retention in MEFs. Differences in Keap1 retention in cells have been interpreted to reflect differences in Keap1 binding to chromatin and to other protein complexes (41, 43).

After culture with vehicle, almost all Keap1 was released from the MEFs during incubations with 0.1% and 0.5% Triton X-100 (Fig. 5A). In contrast, after culture with BIX01294, most of Keap1 was retained in the MEFs during incubations with 0.1% and 0.5% Triton X-100. One fourth of Keap1 was retained in the latter MEFs during incubation with 1% SDS. Most of Keap1 and histone H3 were released together from MEFs that were cultured with BIX01294 (Fig. 5A). Culture with BIX01294 did not affect cJun, lamin B1 or histone H3 retention in MEFs during incubations with the detergents that were tested (Fig. 5A). The stabilization of Keap1 retention in MEFs that were cultured with BIX01294 suggests that BIX01294 strengthened Keap1 interactions in MEFs

**Figure 5.**
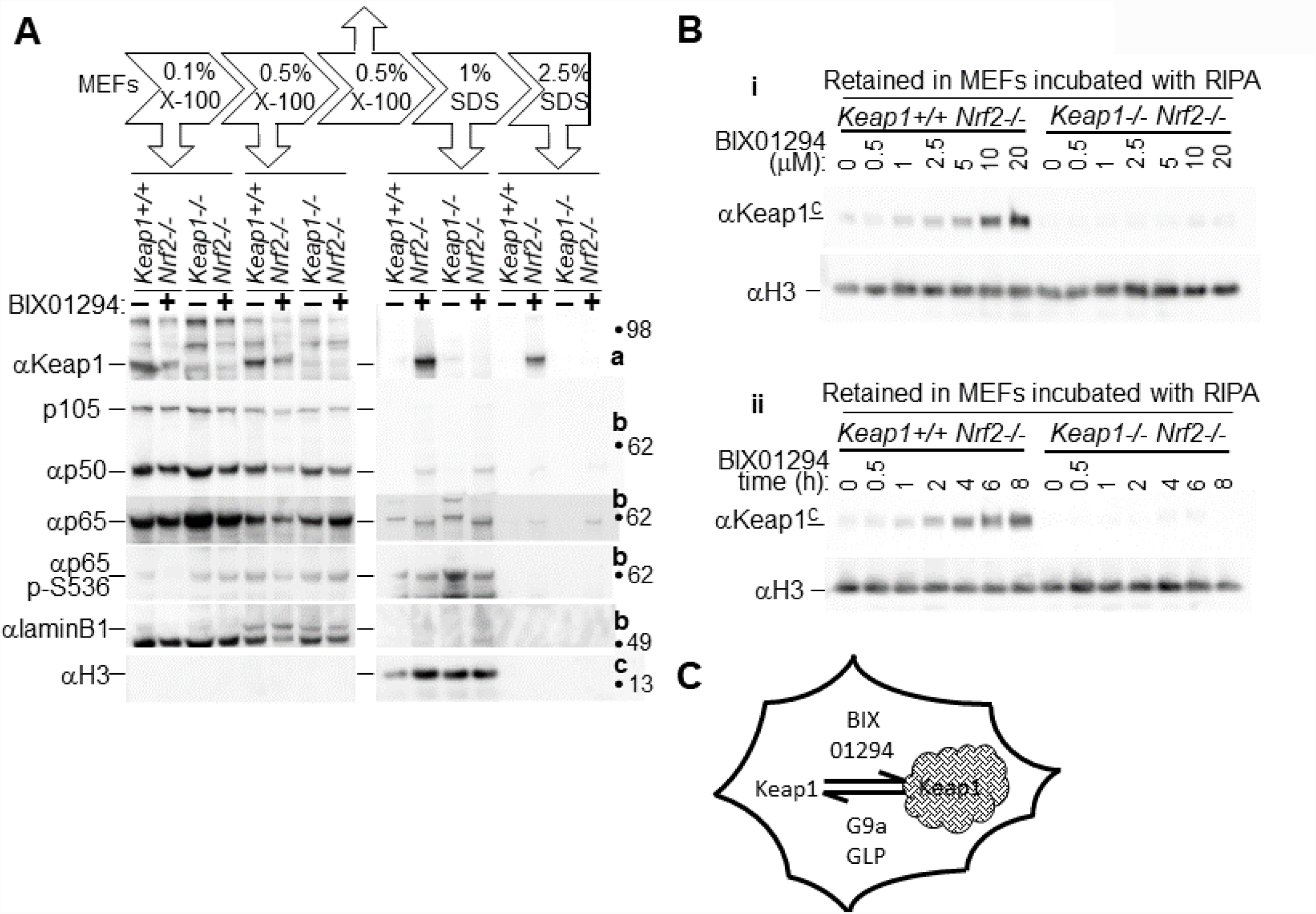
G9a-GLP inhibitors stabilize Keap1 retention in permeabilized MEFs. (A) Culture with BIX01294 stabilizes Keap1 and NFкB p50 retention in MEFs that are incubated with detergents. The MEFs that are indicated above the lanes were cultured with vehicle (-) or with 20 µM BIX01294 (+) for 48 hours. After culture, the MEFs were incubated sequentially in buffers that contained the detergents indicated above the lanes (see Materials and Methods). The proteins that were released during incubation with each detergent were analyzed by immunoblotting using the antibodies indicated to the left of the images. No proteins were detected in the lanes that were loaded with supernatants from a second incubation with 0.5% Triton X-100 (up arrow in the diagram at the top), and those lanes are not shown. The same samples were analyzed on several membranes as indicated to the right of the images (a, b, c). The images show results from a representative experiment in which *Keap1+/+ Nrf2-/-* #2 and *Keap1-/- Nrf2-/-* #4 MEFs were compared. The mobilities of molecular weight markers are indicated to the right of the images. (B) Effects of the concentration of BIX01294 and of the time of culture on Keap1 retention in MEFs. The MEFs that are indicated above the images were cultured with the indicated concentrations (µM) of BIX01294 for 4 h (i, upper panel), and for the indicated times (h) with 20 µM BIX01294 (ii, lower panel). After culture, the MEFs were incubated in RIPA buffer, and the proteins that were retained in the MEFs were analyzed by immunoblotting. The images show αKeap1^C^ and αVinculin immunoblots of the proteins that were retained in the MEFs. The images show the results from a representative experiment in which *Keap1+/+* Nrf2-/- #8 and *Keap1-/-* Nrf2-/- #5 MEFs were compared. (C) The diagram illustrates the stabilization of Keap1 retention in MEFs by G9a-GLP inhibition.

After culture with vehicle, all NFкB p50 was released from the MEFs during incubations with 0.1% and 0.5% Triton X-100 (Fig. 5A). After culture with BIX01294, 10% of NFкB p50 was retained in the MEFs during incubations with 0.1% and 0.5% Triton X-100. BIX01294 did not affect the amount of NFкB p65 that was retained in MEFs. However, S536 phosphorylation increased NFкB p65 retention, and the electrophoretic mobility of retained NFкB p65 was altered by culture with BIX01294 (Fig. 5A).

The concentration of BIX01294 and the time of culture that stabilized Keap1 retention in MEFs were similar to the conditions that augmented Keap1 binding to specific genes and inhibited H3K9me2 deposition (Fig. 5B, 2B). A lower concentration of MS012 than of BIX01294 was required to stabilize Keap1 retention in MEFs as well as to augment Keap1 binding and inhibit H3K9me2 deposition at specific genes (Fig. 3A). BIX01294 stabilized Keap1 retention in both uninfected and in virus infected MEFs, consistent with BIX01294 augmentation of Keap1 binding to different genes in uninfected and in virus infected MEFs.

### Keap1 and electrophiles attenuate the transcription of virus induced genes through opposite effects on different NFкB subunits

We compared Keap1 and electrophile effects on virus induced gene transcription separately and in combination. Virus infection induced higher levels of *Ifnb1* and *Tnf* transcription in *Keap1-/-* MEFs than in MEFs with intact *Keap1*, whether these MEFs were cultured with vehicle or with tBHQ (Fig. 6A). tBHQ reduced and delayed the peak *Ifnb1* and *Tnf* transcript levels to a similar extent both in MEFs with intact *Keap1* and in *Keap1-/-* MEFs. tBHQ reduced viral induction of *Il6* transcription both in *Keap1-/-* MEFs and in MEFs with intact *Keap1*. The combined effects of Keap1 and tBHQ suggest that they moderated virus induced gene transcription independently of each other.

**Figure 6.**
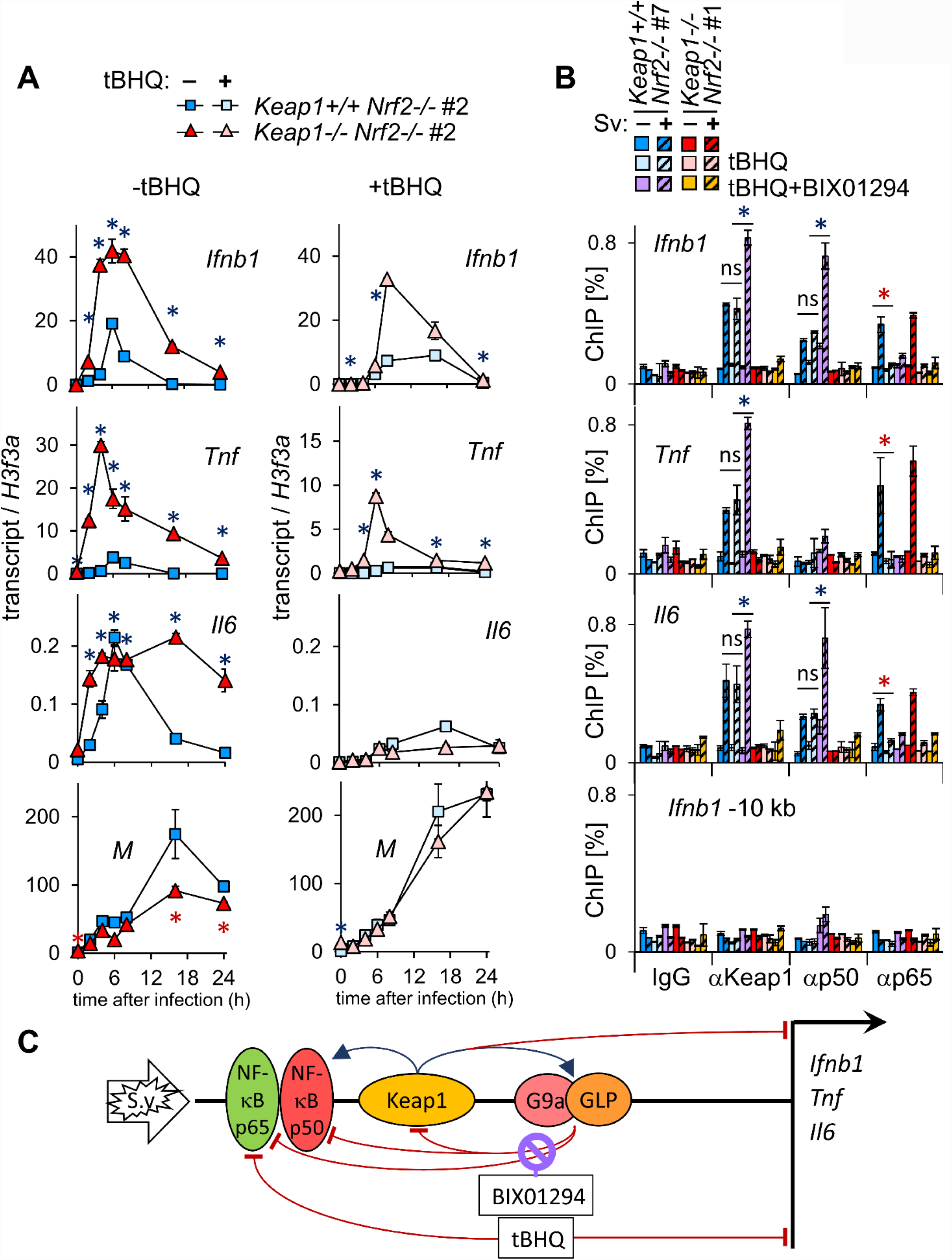
Keap1 and electrophiles attenuate virus induced gene transcription in parallel, and regulate chromatin binding by different NFкB subunits through unrelated mechanisms. (A) Keap1 and tBHQ moderate the transcription of virus induced genes independently of each other. MEFs with intact Keap1 and MEFs with *Keap1-/-* deletions, each with *Nrf2-/-* deletions, were cultured with vehicle (left graphs) or with 50 µM tBHQ (right graphs) for 24 hours before virus infection. The levels of the transcripts indicated in the graphs were measured at the times after virus infection indicated at the bottom. The graphs show the results of a representative experiment in which *Keap1+/+ Nrf2-/-* #2 and *Keap1-/- Nrf2-/-* #2 MEFs were compared. The reproducibility of Keap1 effects on transcription in MEFs that were cultured with tBHQ or with vehicle were evaluated in two independent sets of MEFs by two-factor ANOVA analyses (* p<0.0005, blue: increase, red: decrease). (B) tBHQ inhibits NFкB p65 binding to virus induced genes independently of Keap1. MEFs with intact *Keap1* (cool colors) and MEFs with *Keap1-/-* deletions (warm colors), each with *Nrf2-/-* deletions, were cultured with 50 µM tBHQ, 50 µM tBHQ and 20 µM BIX01294, or vehicle for 24 hours before infection. The levels of Keap1, NFкB p50 and NFкB p65 binding were measured 6 hours after mock (solid bars) or virus (striped bars) infection at the genes indicated in the graphs using the antibodies indicated at the bottom. The graphs show the results of a representative experiment in which *Keap1+/+ Nrf2-/-* #7 and *Keap1-/- Nrf2-/-* #1 MEFs were compared. The reproducibility of tBHQ and BIX01294+tBHQ effects on Keap1, NFкB p50, and NFкB p65 binding were evaluated in two independent sets of MEFs by two-factor ANOVA analyses (* p<0.005, blue: increase, red: decrease). (C) The diagram depicts effects of tBHQ and of BIX01294 on Keap1, NFкB p50 and NFкB p65 binding, on H3K9me2 deposition, and on transcription at virus induced genes. The blue arrows indicate effects that are required for binding or for deposition. The red arcs and bars indicate effects that inhibit or moderate binding, deposition, or transcription.

We examined if tBHQ affected Keap1 binding to virus induced genes. Culture with tBHQ did not alter Keap1 binding to these genes upon virus infection (Fig. 6B). BIX01294 augmented Keap1 binding to *Ifnb1, Tnf* and *Il6* in MEFs that were cultured with tBHQ, suggesting that BIX01294 augmented Keap1 binding to these genes by mechanisms that were unrelated to electrophile responses. The independent effects of Keap1 and of tBHQ on *Ifnb1* and *Tnf* transcription, and the lack of tBHQ effects on Keap1 binding to virus induced genes suggest that Keap1 and tBHQ moderated their transcription through independent mechanisms.

We compared the effects of Keap1 and of tBHQ on NFкB subunit binding to virus induced genes. Keap1 was required for NFкB p50 to bind virus induced genes, whereas tBHQ had no effect on NFкB p50 binding to these genes (Fig. 6B). In contrast, tBHQ inhibited NFкB p65 binding to these genes both in MEFs with intact Keap1 and in *Keap1-/-* MEFs, whereas Keap1 did not affect NFкB p65 binding to these genes in the absence of BIX01294.

Moreover, tBHQ blocked the augmentation of NFкB p65 binding by BIX01294, whereas it had no effect on the augmentation of NFкB p50 binding in the same MEFs. Both Keap1 and tBHQ attenuated virus induced gene transcription and altered the compositions of NFкB complexes that bound to virus induced genes, but the molecular mechanisms for these effects were distinct.

## Discussion

These experiments have identified a regulatory circuit that moderates the transcription virus induced genes. Virus infection actuated this circuit by inducing Keap1 to bind virus induced genes. The parallel actuation of mechanisms that activate and that moderate the transcription of virus induced genes can shape the timing and the amplitude of virus induced gene transcription in response to signals that modulate their expression.

Viral induction of G9a-GLP binding and H3K9me2 deposition at genes whose transcription is induced by virus infection were unexpected. A recent study found that trivalent influenza virus vaccination increases H3K9me2 and other histone modifications that correlate with reduced transcription in CD34+ progenitors and monocyte subsets (44). The increased H3K9me2 correlates with reduced cytokine induction *in vitro* in PBMCs from vaccinated subjects. Vaccination also reduces chromatin accessibility at cytokine, chemokine and other antiviral response genes in classical monocytes. Chromatin accessibility was reduced within one day after vaccination, suggesting that this is an acute response to vaccination. It is possible that the mechanisms whereby Keap1, G9a-GLP and NFкB p50 moderate virus induced gene transcription in MEFs are related to the increased H3K9me2 and reduced chromatin accessibility and cytokine induction in monocytes of vaccinated subjects.

Since Keap1 does not contain a recognized DNA binding domain, it is likely that the contrasting effects of virus infection and of G9a-GLP inhibitors on Keap1 binding to virus induced and to uninduced genes were mediated by interactions with different DNA binding proteins at different genes. It is possible that lysine methylation by G9a-GLP inhibits some of these interactions, which can account for the higher levels and stability of Keap1 binding to chromatin in MEFs that were cultured with G9a-GLP inhibitors. The molecular mechanisms that induced Keap1 to bind at virus induced genes and reduced Keap1 binding at cell cycle associated genes upon virus infection remain unknown.

Viral induction of Keap1 binding to *Ifnb1, Tnf* and *Il6*, and the higher levels of these transcripts in *Keap1-/-* MEFs suggest that Keap1 moderated *Ifnb1, Tnf* and *Il6* transcription in virus infected MEFs by binding to these genes. Keap1 bound to *Cdkn1a* and *Ccng1* in uninfected MEFs, and the levels of these transcripts were higher in uninfected *Keap1-/-* MEFs than in MEFs with intact *Keap1*. Because many mechanisms can influence the basal levels of these transcripts, and because G9a-GLP inhibitors enhanced Keap1 binding to these genes, but had little effect on their transcription, it is not clear if Keap1 binding to these genes affected their transcription in uninfected MEFs.

Keap1 was required for G9a-GLP and NFкB p50 recruitment and for H3K9me2 deposition both within virus induced genes and in regions flanking these genes. The molecular mechanisms whereby Keap1 facilitated G9a-GLP and NFкB p50 recruitment remain unclear. Keap1 forms complexes with NFкB p50 and NFкB p65 that can bind chromatin in living cells, suggesting that they can be recruited in concert. G9a and GLP co-precipitate with NFкB subunits from cell extracts, suggesting that they can also be recruited in concert (22, 45). It is plausible that initial G9a-GLP binding and H3K9me2 deposition within virus induced genes facilitated. subsequent G9a-GLP binding and H3K9me2 deposition in flanking regions. It is possible that changes in chromatin structure mediated the effects of Keap1 and G9a-GLP on NFкB p50 binding. The effects of Keap1 and of G9a-GLP inhibitors on NFкB p50 binding to the *Ccl2* enhancer could also be mediated by DNA looping. This hypothesis is consistent with the observation that both NFкB p50 and Keap1 bind to both the *Ccl2* promoter and enhancer.

The absence of G9a-GLP and NFкB p50 binding, and the higher levels of virus induced gene transcription in *Keap1-/-* MEFs suggest that Keap1 moderated the transcription of virus induced genes by enabling G9a-GLP and NFкB p50 to bind these genes. Because of the interdependent effects of Keap1 and G9a-GLP on chromatin binding by each other and by NFкB p50, it was not possible to distinguish their separate roles in the moderation of virus induced gene transcription. The interdependence of Keap1, G9a-GLP and NFкB p50 binding to, and regulation of, virus induced genes suggest that these proteins are part of a regulatory network whose integrated function moderates the transcription of virus induced genes.

Several structurally and mechanistically dissimilar G9a-GLP inhibitors augmented Keap1 and NFкB p50 binding to virus induced genes and enhanced their transcription. The individual G9a-GLP inhibitors differed in the timing, potency, selectivity and mechanisms of G9a-GLP inhibition. All compounds tested augmented Keap1 and NFкB p50 binding to virus induced genes, and three of the compounds enhanced virus induced gene transcription in parallel with G9a-GLP inhibition. Consequently, G9a-GLP lysine methyltransferase activities moderate both (1) Keap1 binding to virus induced genes, and (2) virus induced gene transcription.

The augmentation of Keap1 and NFкB p50 binding to virus induced genes and the enhancement of their transcription by G9a-GLP inhibitors correlated with the suppression of virus induced H3K9me2 deposition. The causality of these relationships is supported by the interdependent effects of both Keap1 and G9a-GLP on H3K9me2 deposition, on transcription, and on binding by each other at virus induced genes. G9a-GLP inhibitors enhanced transcription and inhibited H3K9me2 deposition only in MEFs with intact *Keap1*, suggesting that G9a-GLP must bind to virus induced genes in order to moderate their transcription. Because both G9a-GLP binding and lysine methyltransferase activity were required for the moderation of Keap1 and NFкB binding to virus induced genes, and of their transcription, it is likely that H3K9me2 deposition moderated both Keap1 and NFкB binding, and transcription at these genes. Other G9a-GLP substrates can also contribute to these effects of G9a-GLP inhibitors.

The lack of correlations between the effects of G9a-GLP inhibitors on H3K9me2 deposition and on Keap1 binding or on transcription at genes that were not induced by virus infection indicates that H3K9me2 deposition is unlikely to moderate Keap1 binding or transcription at these genes. Virus infection had opposite effects on Keap1 binding to virus induced genes and to uninduced genes, consistent with the hypothesis that different mechanisms control Keap1 binding at different classes of genes. The lack of a correlation between H3K9me2 deposition and Keap1 binding at uninduced genes is also consistent with the lack of G9a-GLP binding to these genes in uninfected MEFs. It is likely that lysine methylation of other G9a-GLP substrates affects Keap1 binding at genes that are not induced by virus infection. The differences in the effects of virus infection and of G9a-GLP inhibitors on Keap1 binding to virus induced and uninduced genes are consistent with the differences in their effects on the transcription of these genes.

G9a-GLP inhibitors augmented Keap1 binding to specific genes and stabilized Keap1 retention in permeabilized MEFs with the same potencies and times of exposure. Keap1 retention varies at different stages of the cell cycle, and is consistent with to chromatin binding (41). Docosahexaenoic acid stabilizes Keap1 retention in human primary monocyte-derived macrophages and reduces *CXCL10* and *CXCL11* induction by LPS (43). The effects of G9a-GLP inhibitors on Keap1 binding to specific genes and on Keap1 retention in permeabilized MEFs suggest that both properties of Keap1 are regulated by G9a-GLP lysine methyltransferase activity.

The depletion or inhibition of several other lysine methyltransferases and demethylases influences innate immune response gene transcription. *Jmjd3* demethylase deletion alters the transcription of a subset of LPS induced genes in mouse macrophages by mechanisms that do not correlate with H3K27me3 levels (46). *Kdm2b* deletion and KDM5 inhibitors have opposite effects on the transcription of LPS and interferon response genes in mouse macrophages versus human cancer cells that do not correlate with the H3K4me3 levels at these genes (47, 48). *Ezh2* deletion and inhibitors reduce LPS and poly(I:C) induction of pro-inflammatory genes in mouse macrophages, and they enhance CXCL10 transcription in tumor cell lines through mechanisms that are unrelated to H3K27me2 deposition at these genes (49, 50). The correlation between Keap1 and G9a-GLP effects on H3K9me2 deposition and on the transcription of virus induced genes in MEFs is exceptional among the histone modifications that have been investigated. It is likely that Keap1 influences virus induced transcription also by mechanisms unrelated to G9a-GLP and H3K9me2 deposition. G9a-GLP inhibition had a smaller effect than *Keap1-/-* deletions on viral induction of *Tnf* transcription in MEFs. Nevertheless, the interdependent effects of *Keap1* deletions and of G9a-GLP inhibitors on their binding to virus induced genes, on H3K9me2 deposition, and on transcription in virus infected MEFs are consistent with interdependent effects of Keap1 and of G9a-GLP on H3K9me2 deposition and on transcription at virus induced genes.

The effects of Keap1 mutations in mice and in human cancers have been attributed to changes in Nrf2 activity (2-6, 51). The observations that Keap1 can regulate immunomodulatory gene transcription directly as well as indirectly suggest that the physiological functions of Keap1 can be mediated by many mechanisms (7, 8, 13). Compounds that alter Keap1 moderation of virus induced gene transcription could be beneficial in the prophylaxis and treatment of infectious diseases and immune disorders.

## Acknowledgments

The authors thank Masayuki Yamamoto and Thomas Kensler for mouse strains with the *Keap1-* and the *Nrf2-*deletion alleles. The work was funded by funded by National Institutes of Health National Institute on Drug Abuse (DA030339), which had no role in the design or interpretation of the experiments.

## Materials and Methods

### Experimental Design

MEFs were derived from embryos with *Keap1-/-* and *Nrf2-/-* deletions, or with *Nrf2-/-* deletions alone. These MEFs were infected with Sendai virus or mock infected. The levels of virus induced gene transcripts and of cell cycle associated gene transcripts were compared at different times after virus infection in multiple independent MEFs of each genotype. The *Ifnb1, Tnf* and *Il6* transcripts were chosen for analysis because they were classified in different categories based on previous studies of their activation (52-54). The *Cdkn1a* and *Ccng1* transcripts were chosen for analysis because of the evidence for a role of Keap1 in the cell cycle (41).

Chromatin binding by Keap1 and by other regulators of virus induced gene transcription, and H3K9me2 deposition, were measured by ChIP analyses at virus induced genes and cell cycle associated genes in multiple independent MEFs of each genotype. The binding proteins and H3K9me2 were chyosen for analysis because of previous studies of proteins and histone modifications that correlated with the activation or inhibition of cytokine transcription (22, 23, 53).

The roles of G9a-GLP lysine methyltransferase activities in transcription and in chromatin binding were examined because Keap1 was required for G9a and GLP recruitment to virus induced genes. Multiple independently derived MEFs of each genotype were cultured with G9a-GLP inhibitors prior to virus infection. The G9a-GLP inhibitors that were tested, and the concentrations and times of culture, were selected based on previous studies of compounds that inhibited G9a and/or GLP activities (24-27).

The effects of G9a-GLP inhibitors on the retention of Keap1, NFkB subunits, and other proteins in permeabilized MEFs were tested because of previous findings that Keap1 can be retained in permeabilized cells (41, 43). The effects of tBHQ alone and in combination with *Keap1-/-* deletions on virus induced gene transcription and regulatory protein binding were examined because of previous studies of Keap1 roles in electrophile response gene regulation.

### Derivation of MEFs

*Keap1-/-* and *Nrf2-/-* mice (51, 55) were crossed to generate *Keap1-/+* and *Nrf2-/+* heterozygotes. Primary MEFs were isolated from embryos produced by these mice at about 13 days post coitus. The MEFs were expanded for up to 5 passages and any compounds or vehicle were added to the cultures. MEFs from different embryos are identified in the figures and figure legends by #number after the genotype. All experiments involving live mice were approved by the Institutional Animal Care & Use Committee at University of Michigan.

### MEF culture and Sendai virus infection

The MEFs were cultured in DMEM (GIBCO, cat. no. 11995) supplemented with 10% FBS and 0.1 mg/ml penicillin-streptomycin. MEFs of the indicated genotypes were plated at a density of 200,000/ml. When indicated, compounds or vehicle were added to the MEFs at the times indicated prior to virus infection. The MEFs were infected with Sendai virus by the addition of 200 hemagglutinin units/ml of Sendai virus (Charles River Laboratories, Wilmington, MA) or mock infected. For time course analyses, aliquots of the same Sendai virus stock were thawed and added in an identical manner at different times and the cells were harvested and analyzed in parallel.

### Analysis of transcript levels by RT-qPCR

The MEFs (200,000 plated cells per sample) were washed twice with pre-warmed 1X PBS (Gibco cat. no 10010023), released with 0.05% trypsin-EDTA (Gibco cat no. 25300054), and suspended in culture medium. Total RNA was isolated (RNeasy, QIAgen) and equal amounts of RNA were reverse transcribed (Transcriptor First Strand cDNA synthesis, Roche). The relative amounts of cDNAs corresponding to the indicated transcripts were measured using qPCR. The relative transcript levels were calculated by assuming that they were inversely proportional to 2^Ct^, where Ct equals the number of cycles required to reach threshold fluorescence. The amounts of transcripts in each sample were normalized by the amount of *H3f3a* transcripts in the same sample. The average normalized transcript level for duplicate reactions was plotted with upper and lower error bars corresponding to 2*SD. The oligonucleotide primers for RT-qPCR were obtained from IDT and validated by melt-curve analyses, and had the following sequences:

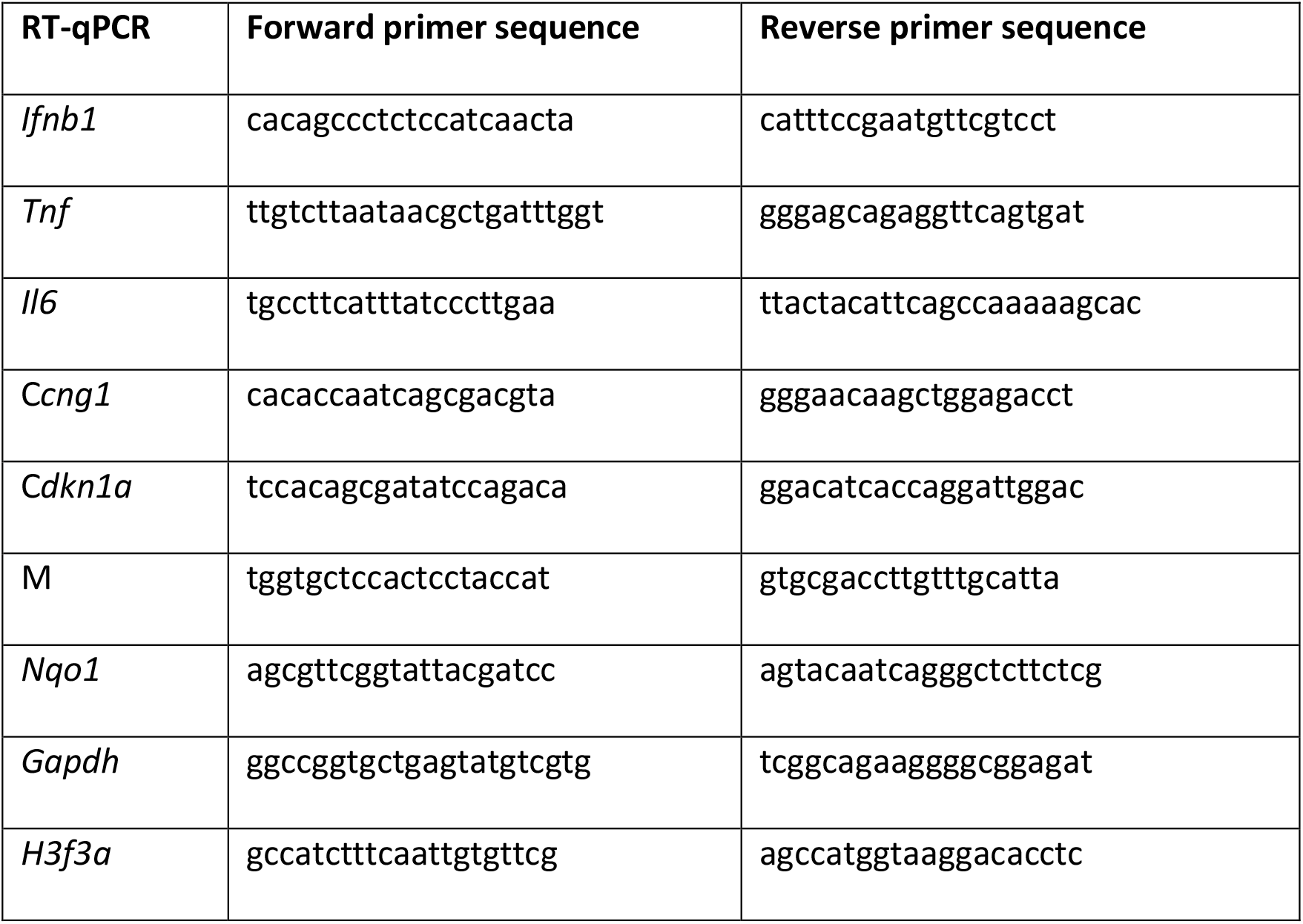

### Analysis of protein binding to specific chromatin regions by ChIP-qPCR

The MEFs (2-8 × 10^7^ cells plated for each genotype and condition) were harvested as described above for transcript analysis. The cells were washed with PBS and crosslinked for 30 min in 2 ml ice cold 1% formaldehyde containing 2X Complete EDTA-free Protease Inhibitor Cocktail (Roche). Cross-linking was stopped by the addition of glycine to 333 mM. The MEFs were collected and lysed by the addition of 4 ml ice cold cell lysis buffer (5 mM PIPES pH 8.0, 85 mM KCl, 0.5% NP-40, and 2X Complete EDTA-free Protease Inhibitor Cocktail (Roche). The nuclei were collected and lysed by 30 min incubation in 4 ml RIPA buffer containing 1X PBS, 1% NP-40, 0.5% sodium deoxycholate, 0.1% SDS, 20 mM N-ethylmaleimide, and 2X Complete EDTA-free Protease Inhibitor Cocktail (Roche). The chromatin was sheared by sonication to produce DNA fragments with a size distribution between 500 bp and 1 kb, followed by 14,000 rpm centrifugation at 4 C for 15 min.

The chromatin was divided into 0.5-1 ml aliquots for immunoprecipitation. Each antibody (2-4 µg per reaction) and IgG control were incubated with Dynabeads (Invitrogen) for at least 4 h at 4C with rotation, washed and resuspended in PBS with BSA. The beads were rotated with the cleared lysates for 24 h at 4 C.

The beads with bound immune complexes were washed 5 times in ice cold 100 mM Tris pH 7.5, 500 mM LiCl, 1% NP-40, 1% sodium deoxycholate, and once with ice cold 10 mM Tris-HCl pH 7.5, 0.1 mM EDTA. Chromatin complexes were eluted by incubating the beads in 1% SDS, 0.1 M NaHCO3 for 1 h at 65 C. The eluted chromatin complexes as well as aliquots of the input chromatin were incubated for 16-20 h at 65 C to reverse the crosslinks. The DNA was purified (QIAquick PCR cleanup, Qiagen). The relative amounts of the genes that were precipitated by each antibody were measured by qPCR. The average percentage of the input DNA that was precipitated (ChIP Efficiency) was plotted for duplicate qPCR reactions with upper and lower error bars corresponding to 2 times the standard deviation. The oligonucleotide primers for ChIP-qPCR were obtained from IDT, were validated by melt-curve analyses, and had the following sequences:

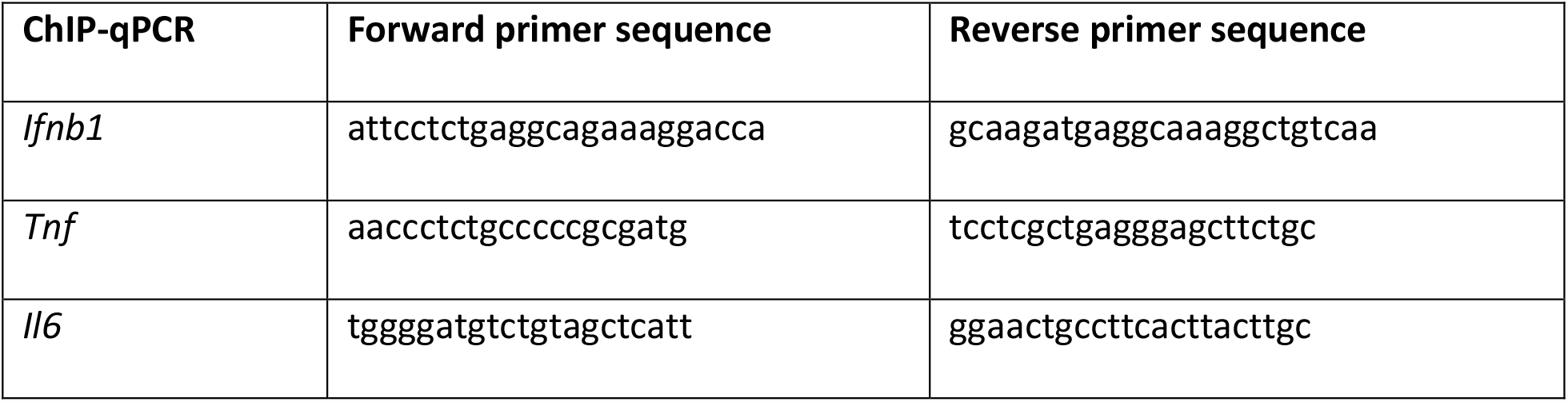
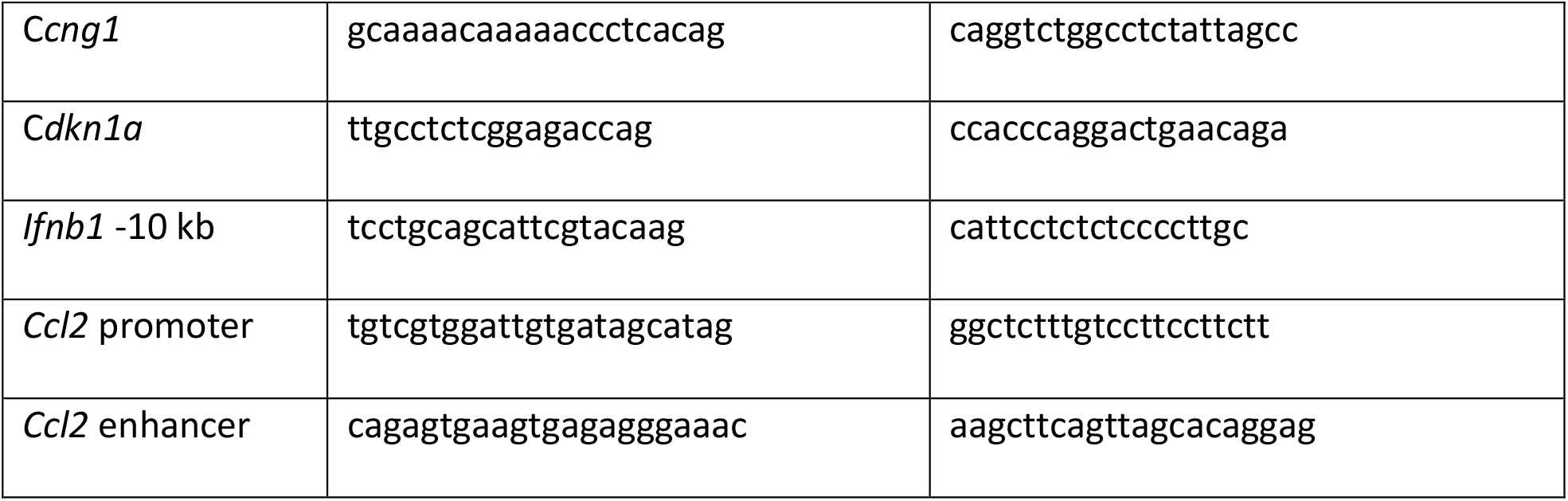

### Analyses of protein retention in MEFs

The MEFs (200,000 plated cells per sample) were cultured as described above and G9a-GLP inhibitor or vehicle was added at the times indicated. Two different protocols were used to evaluate protein retention in MEFs:

#### MEF incubation in buffers with successively stronger detergents

The MEFs were permeabilized by resuspension in 0.2 ml of ice cold CSK buffer containing 0.1% triton X-100, 300 mM sucrose, 100 mM NaCl, 10 mM imidazole pH 7, and 2X Complete EDTA-free Protease Inhibitor Cocktail (Roche) for 30 min on ice. The MEFs were collected by 5 min centrifugation at 4 C. The MEFs were resuspended in in 50 µl ice cold CSK buffer with 0.5% triton X-100, for 30 min on ice. The MEFs were collected by 5 min centrifugation at 4 C. The MEFs were suspended in 50 µl CSK buffer with 1% SDS, for 30 min at RT. The MEFs were collected by 5 min centrifugation at RT. Each of the supernatants and the final pellet were adjusted to 1X SDS loading buffer (58.33 mM Tris·Cl, 5% glycerol, 2.5% SDS, 0.1 M DTT, 0.002% bromophenol blue, pH 6.8) and boiled for 10 min. The proteins were analyzed by polyacrylamide gel electrophoresis and immunoblotting. *MEF incubation in RIPA buffer*. The MEFs were permeabilized by resuspending them in 50 µl ice cold RIPA buffer containing 1X PBS, 1% NP-40, 0.5% sodium deoxycholate, 0.1% SDS, 20 mM N-ethylmaleimide, and 2X Complete EDTA-free Protease Inhibitor Cocktail (Roche) for 30 min on ice. The MEFs were collected by 5 min centrifugation at 4 C. The supernatant and pellet were adjusted to 1X SDS loading buffer, boiled for 10 min, and analyzed by polyacrylamide gel electrophoresis and immunoblotting.

### Antibodies

The αKeap1 (Aviva Systems Biology OACD04962, lot A20190909988) antibody was raised using recombinant Keap1 Met1-Gln286 for immunization. The αKeap1^N^ (Santa Cruz sc-15246, lot H2603) antibody has an epitope mapping near the N-terminus of human Keap1. The αKeap1^C^ (Proteintech 10503-2-AP, lot 00051883) antibody was raised using a recombinant human Keap1 325-624 fragment for immunization The specificities of the three αKeap1 antibodies were evaluated by immunoblotting and by performing parallel RT-qPCR as well as ChiP-qPCR experiments using MEFs with *Keap1-/-* and intact Keap1 alleles. The sources of the other antibodies were as follows: αp50/p105 (Cell Signaling 13586, D4P4d lot 2), αp65 (Cell Signaling 8242, lot 9), αphospho-p65-Ser536 (Cell Signaling 3031, lot 11), αG9a (Cell Signaling 3306, lot 2), αGLP (Abcam ab41969, lot GR191907-41), αH3K9me2 (Abcam ab1220, lots GR3228498-2, GR325223-3), αH3 (Abcam ab1791, lot GR3198255-1), αIRF3 (Santa Cruz sc-9082, lot G0910), αJun (Santa Cruz sc-1694, lot E0207), αvinculin (Cell Signaling 4650T, lot 4), αlaminB1 (Cell Signaling 13435, lot 2), normal rabbit IgG (Santa Cruz sc-2027, lot L2414), normal mouse IgG (Santa Cruz sc-2025, lot I2208).

### Cell culture supplies and chemicals

DMEM (Gibco 11995) was supplemented with 10% (v/v) fetal bovine serum (Atlanta Biologicals S11550 lot H1030) and 0.1 mg/ml penicillin-streptomycin (Gibco 15140). PBS (10010) and 0.05% trypsin-EDTA solution (25300) were from Gibco. The following compounds were obtained from Cayman Chemical: BIX01294 (13124, ≥98%), BRD4770 (11787 ≥98%), UNC0638 (10734, ≥98%), UNC0642 (14604, ≥95%). The following chemicals were obtained from Millipore Sigma: tBHQ (112941, ≥98%) and MS012 (SML2174 ≥98%). These compounds were dissolved in ddH2O (BIX01294) or DMSO (all other compounds) at concentrations ranging from 20 to 50 mM. The final concentration of DMSO in experiments using this vehicle was 0.1%. Other chemicals and detergents were purchased from Millipore Sigma and Thermo Fisher Scientific.

### Statistical analyses of the likelihood of chance occurrence of observed results

The data that are shown in each figure are representative of data that were obtained from multiple independent experiments that were performed using different sets of MEFs (numbers of independent MEFs are specified in the Figure legends). Each graph shows the means and two times the standard deviation for replicate measurements from one experiment. The results of ANOVA analyses to test hypotheses using data from multiple independent experiments with different sets of MEFs are also indicated in each graph. The p-values for each set of experiments were corrected for multiple testing error by the method of Šidák. These p-values were compared to the value indicated in each figure legend to determine if the test supported the acceptance (ns) or rejection (*) of the null hypotheses. The ANOVA tests were performed using the Real Statistics Microsoft Excel plug-in v. 6.8 (Copyright (2013 – 2020) Charles Zaiontz. www.real-statistics.com). ANOVA analyses were performed using data that were obtained from independent experiments.

## Notes

### Competing Interest Statement

The authors have declared no competing interest.

